# Tolerance of combined drought and heat stress is associated with transpiration maintenance and water soluble carbohydrates in wheat grains

**DOI:** 10.1101/2020.05.08.085118

**Authors:** Abdeljalil El Habti, Delphine Fleury, Nathaniel Jewell, Trevor Garnett, Penny J. Tricker

## Abstract

Wheat (*Triticum aestivum* L.) production is increasingly challenged by simultaneous drought and heatwaves. We assessed the effect of both stresses combined on whole plant water use and carbohydrate partitioning in eight bread wheat genotypes that showed contrasting tolerance. Plant water use was monitored throughout growth, and water-soluble carbohydrates (WSC) and starch were measured following a three-day heat treatment during drought. WSC were predominantly allocated to the spike in modern Australian varieties, whereas the stem contained most WSC in older genotypes. Combined drought and heat stress increased WSC partitioning to the spike in older genotypes but not in the modern varieties. Glucose and fructose concentrations in grains measured 12 days after anthesis were associated with final grain weight in the main spike. At the whole plant level, combined drought and heat stress differentially altered daily water use and transpiration response to vapour pressure deficit during grain filling, compared to drought only. Final grain yield was increasingly associated with aboveground biomass and total water use with increasing stress intensity. Ability to maintain transpiration, especially following combined drought and heat stress, appears essential for maintaining wheat productivity.

**One sentence summary:** Higher yield following drought and heat stress in wheats that maintain transpiration and have higher water-soluble carbohydrates content in grains.

## Introduction

Recent decades have witnessed severe drought and heatwaves worldwide, including in major wheat producing regions such as India, the U.S.A., Russia, Western Europe and Australia. These climatic conditions have a significant impact on global wheat production, with dramatic social and economic consequences (van Dijk et al., 2013). Current climate projections predict drought and heatwaves will become more common and more intense in the future (Rosenzweig et al., 2014). One way to limit the impact of weather variability on productivity is to develop wheat varieties better adapted to the changing climate. This can be assisted by understanding the mechanisms underlying plants’ responses to complex stresses so as to identify the traits that characterise stress tolerant varieties for breeding.

A small number of studies document the impact of combined drought and high temperature on wheat productivity and biological processes, especially during the reproductive developmental stage. The effect of combined stresses is more detrimental than the effect of an individual stress (Mittler, 2006). Both drought and high temperatures reduce expansive growth, accelerate flowering and shorten grain filling duration, resulting in a low grain set, size and weight (Asana and Williams, 1965; Pradhan et al., 2012). In combination, drought and high temperature impair the photosynthetic system, reduce stomatal conductance and gas exchange, and disrupt plants’ water relations (Machado and Paulsen, 2001; Shah and Paulsen, 2003). These additive alterations of morphological, physiological and cellular processes result in severe reductions in final grain weight.

Although the major impact of combined drought and high temperature on wheat productivity is well described, there is scarce information on the mechanisms that determine the ability to maintain grain weight in these unfavourable environments (also called tolerance). Wheat harvested grain mass consists of 85 % carbohydrates, of which ∼80 % is starch (Stone and Morell, 2009). During grain filling, water-soluble carbohydrates (WSC) are delivered to grains either from current photosynthesis in photosynthesising organs or from remobilisation of WSC stored during the vegetative stage (Borrell et al., 1989; Schnyder, 1993). Abiotic stress after anthesis can limit gas exchange and damage the photosynthetic system, in which case stored carbohydrates become a major source of carbon for grain filling (Blum, 1998). In addition to the contribution from stem reserves, spike organs, especially awns, are thought to contribute to the grain filling process due to their active photosynthesis, especially in dry environments (Evans et al., 1972; Rebetzke et al., 2016).

Carbohydrate synthesis and transport are closely related to water movements in plants. Open stomata are necessary for carbon capture and plants trade-off between maximising carbon assimilation and limiting water loss through transpiration under adverse conditions such as drought. Carbohydrate transport via the phloem and distribution throughout the plant rely on water exchange with adjacent xylem (Hölttä et al., 2009), and the impact of water shortage on xylem water transport also impairs phloem function (Sevanto, 2018). Soluble carbohydrates also play an important role during drought by acting as compatible osmolytes to maintain cell turgor and favourable plant water status, thereby sustaining biological processes and soil water uptake (Blum, 2017). Maintaining plant hydration and enhancing carbohydrate remobilisation to grains are considered key factors for crop productivity in limiting environments (Blum, 2006), and the interplay between plant water relations and carbohydrate metabolism and distribution highlights the importance of studying both mechanisms together. In this work, we describe the impact of combined drought and heat stress on whole plant water use and carbohydrate partitioning during grain filling in diverse wheat genotypes. We hypothesised that the combination of both stresses would alter plant water use and carbohydrate partitioning in the stem and spike, and that WSC availability would be a limiting factor for optimal grain weight under combined drought and heat stress.

## Results

### Combined drought and heat stress differentially reduced total grain weight at harvest

The effect of drought and combined drought and heat stress (D&H) on total grain weight per plant at harvest (per plant yield) depended on genotype (Table 1). Drought reduced per plant yield in Currawa, Odessa, Frame, Young and Gladius (Fig. 1). Interaction of drought with high temperature further reduced per plant yield in Odessa, Mendos and Young. In contrast, heat stress did not exacerbate the effect of drought on yield in Currawa, Frame and Gladius. Per plant yield in Koda was not sensitive to either drought or D&H stress. Overall, the combination of drought and high temperature was more detrimental to per plant yield in some genotypes, but not all (Fig. 1a).

**Table 1.** Analysis of variance (ANOVA) showing the statistical significance of the traits measured for genotype, treatment and interaction between genotype and treatment.

| Traits | Genotype | Treatment | Interaction |
| --- | --- | --- | --- |
| Total grain weight per plant | *** | *** | ** |
| Total grain weight in the main spike at 12 DAA | *** | ns | ns |
| Total grain weight in the main spike at harvest | *** | *** | ns |
| Aboveground biomass (excluding grains) | *** | *** | ns |
| Plant height | *** | ns | ns |
| Total water use | *** | *** | ns |
| Water use after treatment | ** | ** | ns |
| biomass WUE | *** | *** | * |
| grain WUE | *** | *** | *** |
| Harvest index | *** | *** | *** |
| WSC concentration in grains | *** | *** | ** |
| Starch concentration in grains | *** | *** | *** |
| Total carbohydrates concentration in grains | *** | ** | * |

\* $p < 0.1$ ; \*\* $p < 0.01$ ; \*\*\* $p < 0.001$ ; ns not significant

**Figure 1.**
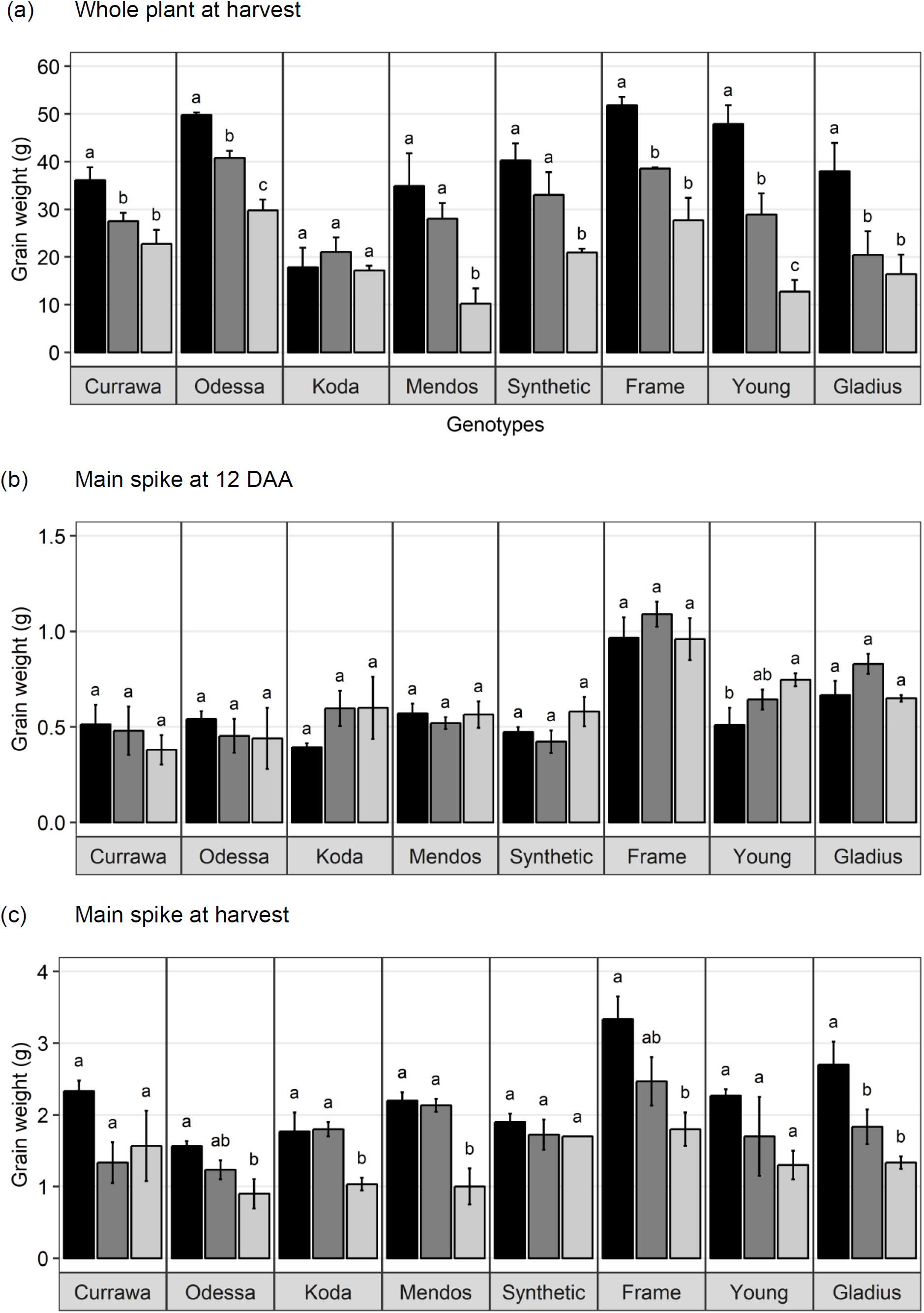
Combined drought and heat stress differentially reduced final grain weight. (a) Mean total grain weight per plant at harvest (n=4). (b) Mean total grain weight in the main spike at 12 days after anthesis (n=3). (c) Mean total grain weight in the main spike at harvest (n=4). Error bars are standard error. Letters indicate the results of Tukey’s test comparing treatment effect within each genotype (p < 0.1). Plants were grown under well-watered conditions (black), drought (dark grey) or combined drought and heat (light grey).

In order to assess the impact of drought and combined D&H on grain filling, grain dry weight in the main spike was measured at 12 days after anthesis (DAA) and at harvest. At 12 DAA, total grain weight in the main spike was different among genotypes (Fig. 1b) but there was no effect of the treatments or genotype x treatment interaction (Table 1). At harvest, total grain weight in the main spike was reduced by drought in Gladius, and reduced by combined D&H in Odessa, Koda, Mendos and Frame (Fig. 1c). There was no effect of drought or combined D&H on the main spike total grain weight at harvest in Currawa, Synthetic W7984 and Young.

### The correlation between aboveground biomass, water use and plant yield increased with increasing stress intensity

Plant yield was associated with aboveground vegetative biomass and total water use to different degrees depending on treatments (Fig. 2). Total grain weight was linearly related to aboveground biomass and total water used by the plant throughout the experiment; these coefficients increased with stress intensity from r^2^ =0.2 and r^2^ =0.31 in well-watered conditions (WW), to r^2^ =0.35 and r^2^ =0.56 under drought, and r^2^ =0.46 and r^2^ =0.67 under combined D&H, respectively (Fig. 2a). This dependence of plant yield on total water use and interaction with treatment was confirmed in a repeated experiment (Supp. Fig. S1). During the 3d heat treatment, plants generally used similar amounts of water as compared to well-watered conditions (Supp. Fig. S2), although soil water potential was halved.

**Figure 2.**
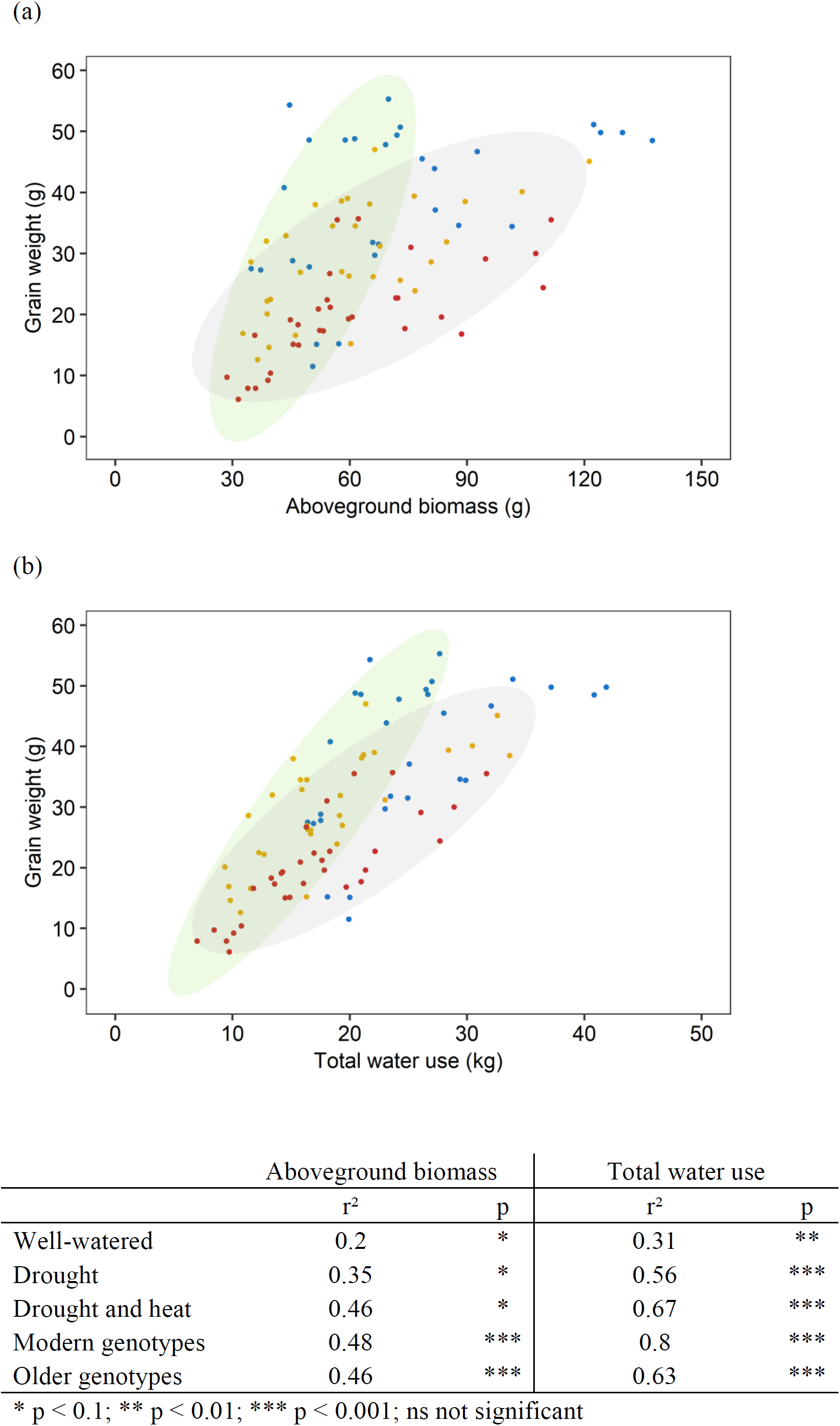
Aboveground biomass and water use explained more of the variation in grain weight under increasing stress intensity than in well-watered conditions. Relationships between final grain weight per plant and (a) aboveground biomass excluding grains, (b) total water use. Each point represents one plant, grown under well-watered conditions (blue), drought (orange) or combined drought and heat stress (red). Ellipses circle modern genotypes (in green) and older genotypes (in grey). Table shows r^2^ and p-value of linear regressions (‘*’ p < 0.1, ‘**’ p < 0.01, ‘***’ p < 0.001) for each treatment (well-watered, drought, combined drought and heat stress) and genotype group (modern, older).

When comparing modern Australian genotypes (Frame, Young and Gladius) to older genotypes (Currawa, Odessa, Koda, Mendos and Synthetic W7984), the relationship between per plant yield and aboveground biomass was similar in both groups (r^2^=0.48 and r^2^=0.46, p < 0.001, respectively). Despite this, the slope of the regression for modern varieties was higher compared to older genotypes (a=0.8 and a=0.3, respectively, Fig. 2). The higher slope for modern genotypes was explained by the lower biomass required to produce similar grain weight compared to older genotypes under well-watered conditions and reflects the high harvest index of modern genotypes in favourable conditions. However, the dependence of plant yield on total water used was higher in modern genotypes compared to older genotypes (r^2^ =0.80 and r^2^ =0.63, p < 0.001, respectively).

### Combined drought and heat stress differentially reduced transpiration response to vapour pressure deficit

As total water use was strongly dependent on plant biomass (Supp. Fig. S3), water use was normalised to the final aboveground biomass and expressed as unit of water per unit of biomass to allow for comparison between plants. Water use differed between genotypes following combined D&H when all treated plants (D and D&H replicates) were in the same droughted conditions (Fig. 3a). Interaction of drought and 3d high temperature reduced daily water use in Odessa, Koda, Mendos and Young for the subsequent 30d, whereas daily water use following combined D&H was similar to D alone in Currawa, Frame, Synthetic W7984 and Gladius.

**Figure 3.**
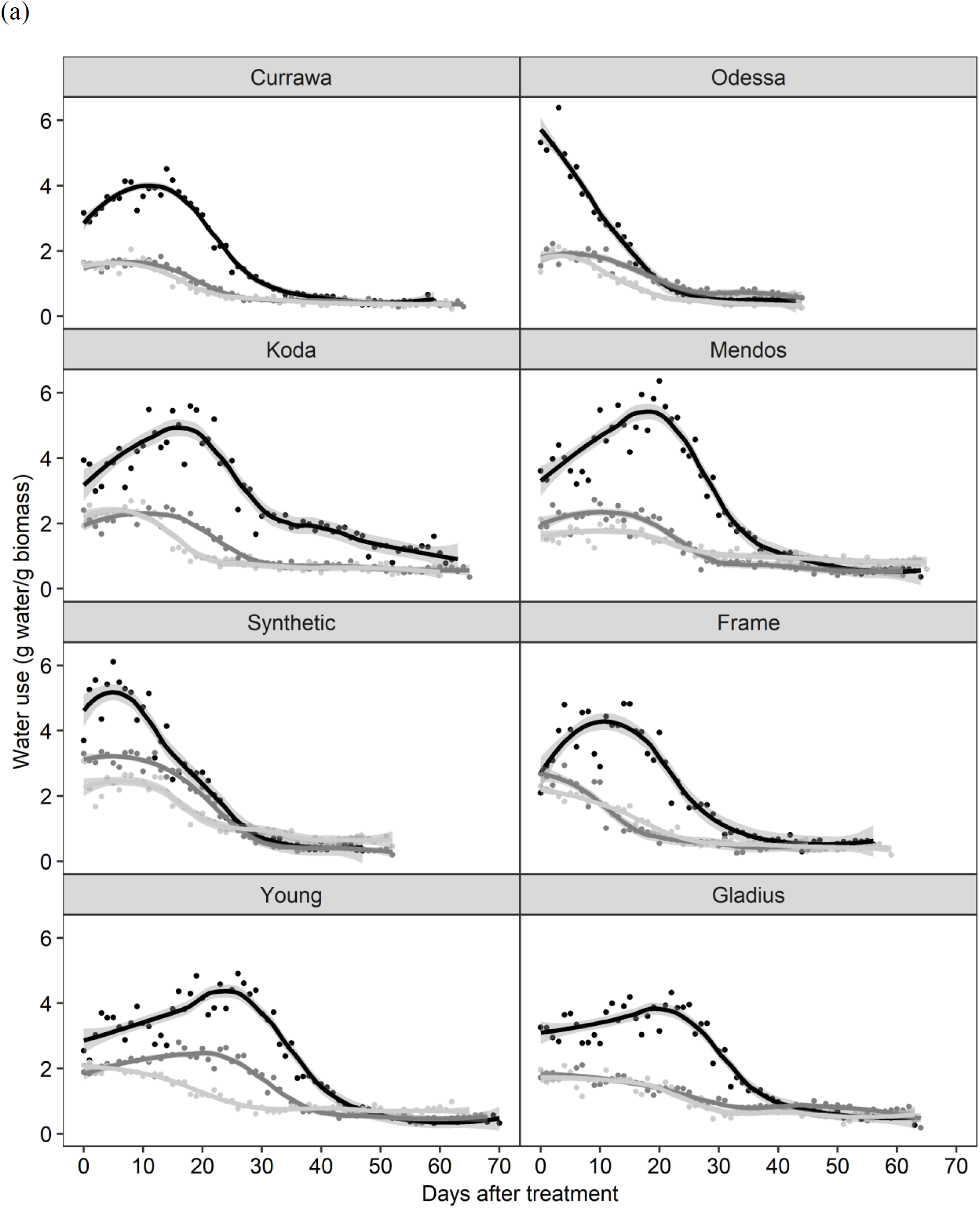
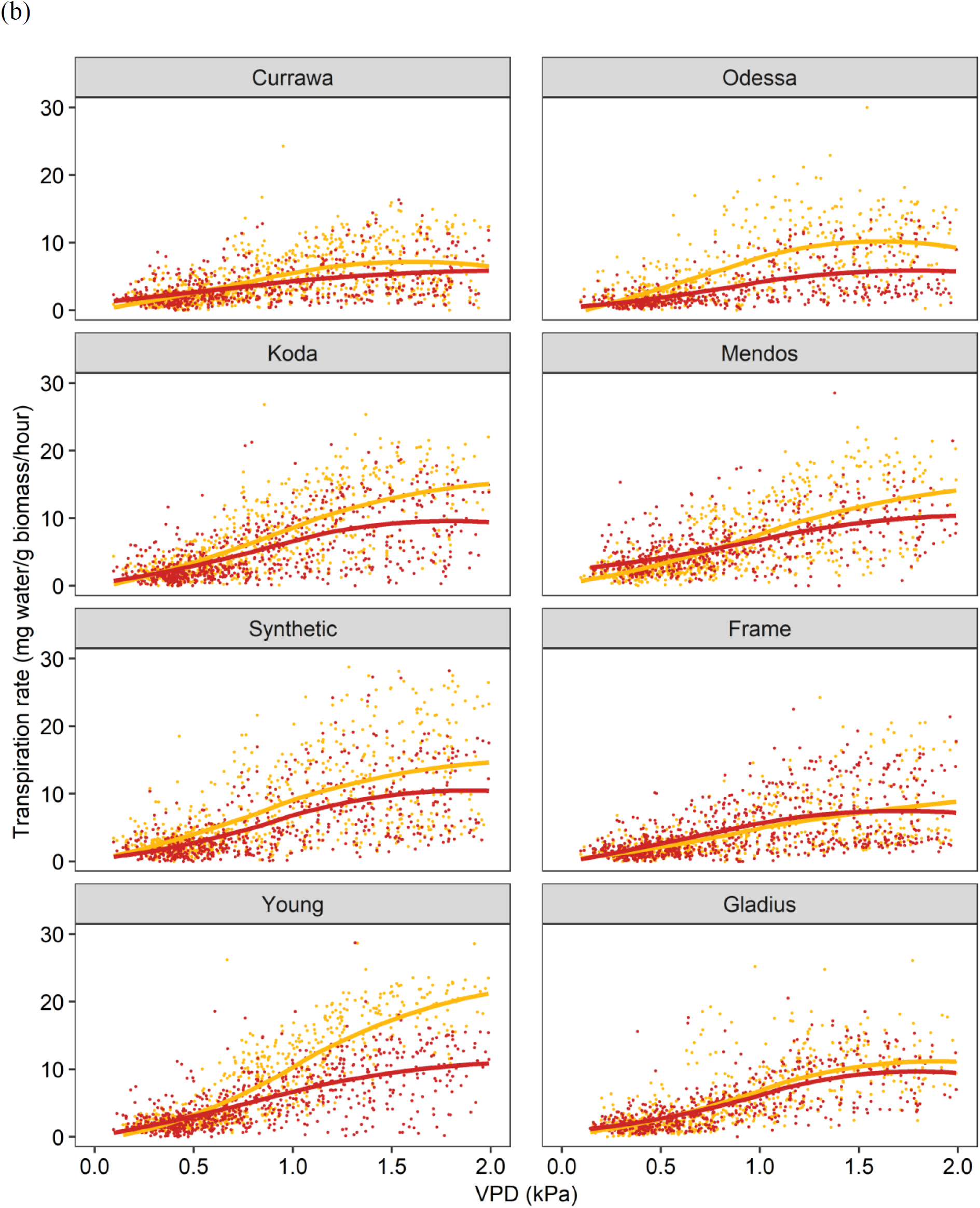
Interaction of high temperature and drought differentially reduced daily water use and transpiration response to vapour pressure deficit. (a) Daily water use per plant estimated as total irrigation per day, normalised to aboveground biomass. Plants grown in well-watered conditions (black), drought (dark grey) or in drought following three-day heat stress (light grey). 0 DAT is the first day post heat treatment (12 days after anthesis). Trend lines are loess regressions. Values are means of four replicates (n=4). The confidence interval (0.95) is displayed around smoothed regressions in grey. (b) Hourly transpiration rate response to VPD normalised to aboveground biomass. Plants grown in drought (orange) or in drought following three-day heat stress (red). Graphs include data from 0 DAT to 30 DAT. Trend lines are smooth regression lines.

Plant water use is the summation of transpiration, which is driven by changes in vapour pressure deficit (VPD). Transpiration response to VPD was differentially altered by the three-day high temperature treatment depending on genotype over the same grain-filling period when D and D&H plants were in the same droughted conditions (Fig. 3b). For both transpiration rate (TR) and specific transpiration rate (STR), statistical significance was confirmed (p < 0.05) for the genotype × treatment interaction component of the VPD slope parameter. That is, the VPD effect on transpiration exhibited genotype × treatment interaction both before and after normalisation to final aboveground biomass.

Heat stress reduced subsequent transpiration rate at VPD > 0.5 kPa in Odessa, at VPD > 0.7 kPa in Young and at VPD > 1.0 kPa in Koda and Mendos (Fig. 3b). In contrast, transpiration response to VPD was not altered following combined D&H stress in Currawa, Frame, Synthetic W7984 and Gladius. Transpiration response to VPD was affected in the same genotypes where daily water use was reduced by combined D&H. During drought, Young had the highest transpiration rate at VPD > 1.5 kPa whereas Currawa and Frame had the lowest transpiration rates. Following combined D&H, Synthetic W7984 had the highest transpiration rate at VPD > 1.5 kPa whereas Currawa and Odessa had the lowest transpiration rates (Supp. Fig. S4b).

### Combined drought and heat stress increased WSC partitioning to the spike in old genotypes, but not in modern varieties

WSC were quantified in the main stem and the spike tissues following all treatments 12 DAA, i.e. immediately following the heat stress for the D&H replicates. There was a clear contrast between older genotypes (Currawa, Odessa, Koda, Mendos, Synthetic W7984) and more modern varieties (Frame, Young, Gladius) for WSC partitioning in stem parts compared with the spike (Fig. 4). In well-watered conditions, stem parts contained 67-87 % of total WSC in older wheat genotypes compared to 28-50 % in modern varieties. The spike tissues (excluding grains) contained 49-71 % of total WSC in modern varieties, whereas WSC in the spike were 12-33 % of total WSC in older genotypes.

**Figure 4.**
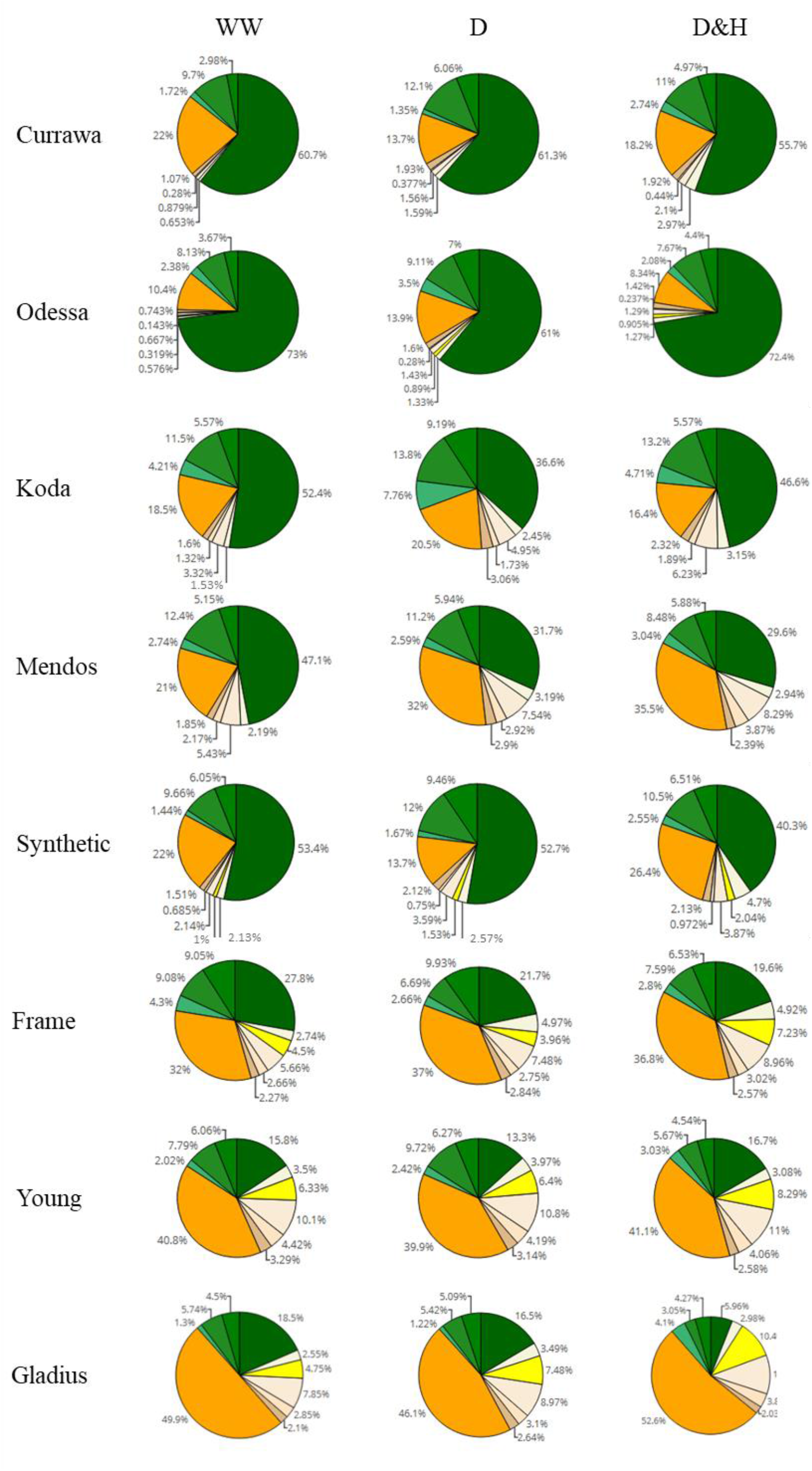
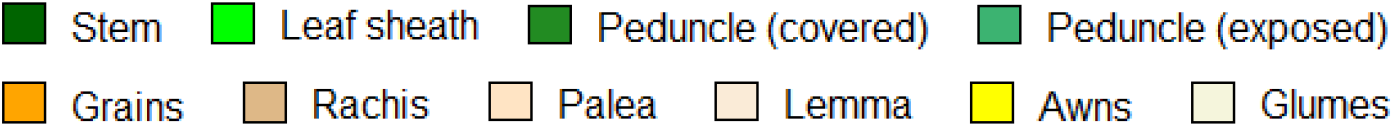
Combined drought and heat stress increased WSC partitioning to the spike in old genotypes, not in modern varieties. Genotypes are shown in order of date of release (top – bottom, oldest = Currawa to newest = Gladius). Total water-soluble carbohydrates (WSC) as in different organs of plants as a percentage of total WSC (n=3). Plant organs are colour-coded as shown in the legend.

Drought and combined D&H differentially affected WSC distribution in the main stem and spike depending on genotype (Fig. 4). Drought significantly increased WSC partitioning into the spike in Currawa and Odessa and reduced the WSC fraction in the stem in Frame (Supp. Table 1). Combined D&H significantly increased WSC partitioning into the spike in older genotypes, whereas there was no change in WSC partitioning in modern varieties in both treatments. Changes in WSC allocation to the spike following combined drought and heat stress did not affect WSC partitioning to grains, except in the Synthetic genotype where WSC partitioning to grains was significantly increased by combined D&H compared to drought only.

### The relationships between WSC concentrations and plant yield depended on plant organ, individual carbohydrate and the date of release of the variety

The relationship between WSC concentration at 12 DAA and plant yield depended on tissue and date of variety release. Total WSC concentration in the stem at 12 DAA was positively related with per plant yield in modern genotypes (r^2^ =0.53) whereas there was no significant regression observed in older genotypes (r^2^ =0.1) (Table 2, Fig. 5a). In contrast, total WSC concentration in awns at 12 DAA was positively related with plant yield in the two, awned older genotypes (r^2^ =0.85) whereas there was no relationship in modern genotypes (r^2^ =0.06, Fig. 5b). Similarly, in grains, total WSC concentration at 12 DAA was positively related with plant yield in older genotypes (r^2^ =0.41) whereas there was no relationship in modern genotypes (r^2^ =0.13) (Table 2, Fig. 5c).

**Table 2.** r^2^ and p-values of linear regressions between individual carbohydrates’ concentrations in different plant organs at 12 DAA and final grain weight per plant in modern vs. older genotypes.

| Carbohydrate | Plant tissue | Modern |  | Old |  |
| --- | --- | --- | --- | --- | --- |
| | | $r^2$ | p | $r^2$ | p |
| WSC | Stem | 0.53 | **0.03** | 0.1 | 0.26 |
|  | Awns | 0.06 | 0.54 | 0.85 | **0.01\*** |
|  | Grains | 0.13 | 0.33 | 0.41 | **0.01** |
| Glucose | Stem | 0.34 | **0.1** | 0.01 | 0.73 |
|  | Awns | 0.14 | 0.33 | 0.84 | **0.1** |
|  | Grains | 0.47 | 0.13 | 0.00 | 0.9 |
| Fructose | Stem | 0.08 | 0.45 | 0.31 | **0.03** |
|  | Awns | 0.11 | 0.39 | 0.81 | **0.01** |
|  | Grains | 0.54 | **0.09** | 0.06 | 0.63 |
| Sucrose | Stem | 0.56 | **0.02** | 0.13 | 0.18 |
|  | Awns | 0.07 | 0.49 | 0.67 | **0.05** |
|  | Grains | 0.44 | 0.15 | 0.05 | 0.66 |
| Fructan1 | Stem | 0.17 | 0.27 | 0.1 | 0.25 |
|  | Awns | 0.47 | **0.04** | 0.35 | 0.22 |
|  | Grains | 0.04 | 0.7 | 0.08 | 0.6 |
| Fructan2 | Stem | 0.73 | **0.003** | 0.23 | **0.07** |
|  | Awns | 0.05 | 0.56 | 0.44 | 0.15 |
|  | Grains | 0.47 | 0.13 | 0.24 | 0.32 |
| Starch | Grains | 0.08 | 0.46 | 0.14 | 0.18 |

\* Only two older genotypes were awned.

**Figure 5.**
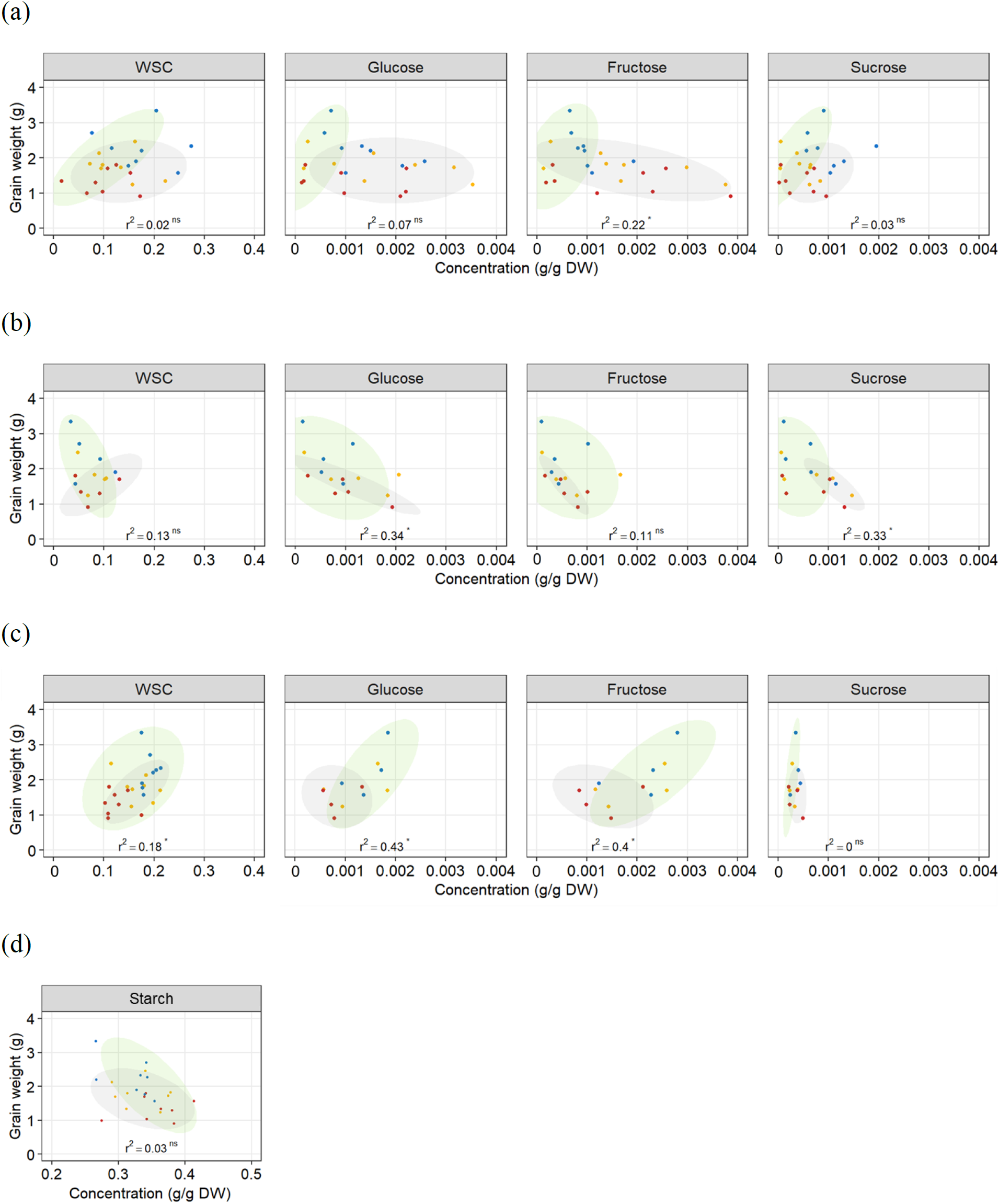
Relationship between total WSC, glucose, fructose, sucrose and starch concentrations in (a) the stem, (b) awns, (c-d) grains at 12 DAA, and final grain weight per plant. Each point represents an average of carbohydrate concentration (n=3) in the main tiller for one genotype and one treatment. Ellipses circle old genotypes (grey) and modern genotypes (green). Plants grown under well-watered conditions (blue), drought (orange) or combined drought and heat stress (red). Regression lines include all datapoints. r^2^ and p-value of linear regressions (‘ns’ not significant, ‘*’ p < 0.05) are indicated.

To determine whether individual WSC varied similarly to total WSC, we quantified glucose, fructose and sucrose concentrations in the stem, awns and grains. In the stem, a similar contrast was observed for individual WSC between older and modern varieties as was observed for total WSC (Table 2, Fig. 5a). Glucose and sucrose concentrations in the stem at 12 DAA were positively related with plant yield in modern varieties (r^2^ =0.34 and 0.56, respectively). Stem fructose concentrations at 12 DAA were negatively related to plant yield in older varieties (r^2^ =0.31). In awns, glucose, fructose and sucrose concentrations at 12 DAA were each negatively related with plant yield in the two awned older genotypes (r^2^ =0.84, 0.81 and 0.67 respectively) but there were no significant relationships between these sugars and plant yield in more modern types (Table 2, Fig. 5b). In grains, unlike other tissues, sucrose concentrations at 12 DAA were low compared to glucose and fructose concentrations (Fig. 5c). Glucose and fructose concentrations were positively related with total grain weight at harvest (r^2^=0.43 and 0.40, respectively – Fig. 5c). Two (unknown) fructans also appeared important for plant yield in modern varieties: fructan 1 in awns (r^2^ =0.47) and fructan 2 in the stem (r^2^ =0.73). In contrast with other sugars in the awns, fructan 1 concentrations at 12 DAA and yield per plant were positively related. There was no relationship between starch concentrations at 12 DAA and plant yield (Fig. 5d).

### Drought and combined drought and heat stress altered WSC and starch balance in grains

In order to quantify WSC availability for starch synthesis, WSC and starch concentrations were measured in grains 12 DAA, immediately following D&H treatments. There was a significant interaction between genotype and treatment for WSC and starch concentrations in grains (Table 1). Drought significantly reduced WSC concentration in grains in the Synthetic type and heat stress did not exacerbate the effect of drought (Fig. 6). WSC concentration in grains was reduced by combined D&H in Odessa and Koda compared to WW and was specifically reduced by combined D&H in Currawa, Young and Gladius. Starch concentrations offset the reduction in WSC concentration in Currawa, Odessa and Synthetic W7984, resulting in a similar total non-structural carbohydrates (NSC) concentration in grains in all conditions. The balance between WSC and starch concentrations was altered in Koda, Young and Gladius (Fig. 6). Total NSC concentration was reduced under drought in Koda and Young, and combined D&H reduced total NSC in Gladius. There was no significant effect of drought and combined D&H on WSC and total NSC concentration in grains in Currawa, Odessa, Mendos and Frame. Overall, there was a significant interaction between genotypes and treatments for total carbohydrates concentration in grains (Table 1) that was mainly driven by interaction between genotypes and treatments for WSC concentration.

**Figure 6.**
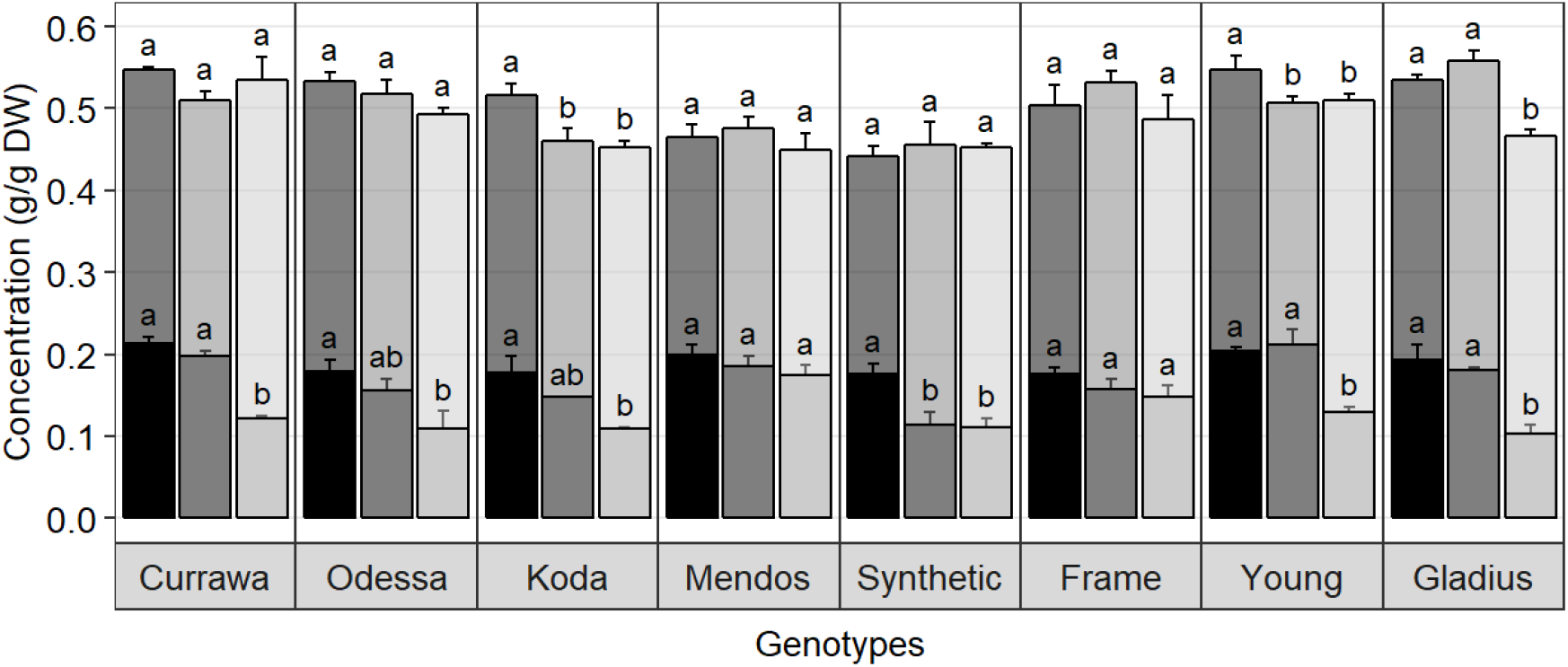
Drought and combined drought and heat stress altered WSC and starch balance in grains at 12 DAA. Water-soluble carbohydrates (solid) and starch (transparent) concentration in grains at 12 DAA from plants grown under well-watered conditions (black), drought (dark grey) or combined drought and heat stress (light grey). The sum of WSC and starch concentrations constitutes the non-structural carbohydrate (NSC). Values are means of three replicates (+/- SE). Letters (top: NSC, bottom: WSC) indicate the results of Tukey’s test comparing treatment effects within each genotype (p < 0.1).

## Discussion

### Higher water use and responsiveness to evaporative demand are indicators of higher yield under combined drought and heat stress

In this study, the impact of drought and combined D&H on total grain weight at harvest was assessed in eight diverse wheat genotypes released between 1912 and 2007. The detrimental effect of heat stress combined with drought depended on genotype, illustrating genetic variation in grain weight response to combined D&H in wheat.

Per plant yield was increasingly dependent on both aboveground biomass and total water use with increasing stress intensity (drought, then additional heat stress) (Fig. 2), highlighting the important relationships between biomass, water use and grain weight under stress previously reported (Reynolds et al., 2006; Blum, 2009; Reynolds and Langridge, 2016). Biomass and water use are linearly related (de Wit, 1958) and mutually dependent during the plant’s lifecycle. During the vegetative stage, transpiration drives biomass accumulation, which in turn results in high water use during grain filling when water is available. The maintenance of water use ensures the favourable water status of plant tissues and assimilate transport to grains.

We observed a high association between per plant yield and total water used in two independent experiments (Fig. 2b, Supp. Fig. S1) regardless of transpiration sensitivity to heat stress: the higher the water use, the higher the total grain weight at harvest. This indicated that maintaining transpiration following heat stress was a desirable trait in our conditions, confirming the strong relationship between plant transpiration and yield (de Wit, 1958; Fischer and Turner, 1978; Sinclair et al., 2005).

The amount of water used in transpiration is driven by the evaporative demand in the atmosphere (Grantz, 1990). Our work illustrated the genetic variation in transpiration response to VPD previously observed in diverse wheat genotypes grown under well-watered and water-limited conditions (Schoppach and Sadok, 2012; Schoppach et al., 2016; Medina et al., 2019). In addition, we identified genetic variation in transpiration response to combined D&H (Fig. 3b). A three-day heat treatment altered transpiration response to VPD in the subsequent drought-only treatment in some genotypes.

As transpiration largely depends on green leaf area, the dynamics of senescence in response to heat stress could potentially have explained genotypic differences in transpiration response to heat stress; water use would be quickly reduced in genotypes with faster heat-induced senescence compared to genotypes with slower senescence rate following heat stress. However, no significant differences or genetic variation for drought and heat stress-induced chlorophyll content (greenness) in comparison with drought were found in these genotypes in repeated experiments (Schmidt et al., 2020). The combination of high evaporative demand and water scarcity can lead to the disruption of the water column in the xylem and cause cavitation. Cavitation damage might explain the lack of recovery in water use following heat stress observed in some genotypes in our experiments.

Here, with both drought and combined D&H stress, grain water use efficiency (WUE) was more important than WUE *per se*. In our experiments, we measured the effects of stress during grain filling when vegetative biomass was already accumulated, rather than during vegetative biomass production. In the field, where increased rooting depth to access available water and early vigour to enhance canopy coverage and reduce soil evapotranspiration are important, this might not be the case. Nonetheless, the transpiration driven water use changes observed here following heat stress during grain-filling, when plants were all subjected to drought, influenced final grain weight. This trait – ability to maintain transpiration following heat stress under drought - separated more from less tolerant types.

Although studied in a limited number of genotypes, we observed clear increased grain weight per unit of biomass and per unit of water in more modern compared with older genotypes. In addition to improved plant architecture for assimilation and partitioning (harvest index), we also observed striking differences in the partitioning of assimilates between vegetative and spike tissues before the imposition of stress.

### The spike is the main storage tissue for WSC in the more modern wheat varieties

Excess assimilates that are not used for growth and defence may be stored for further use during reproductive stages. The stem is considered an important source of stored WSC for grain filling, and the ability to store and remobilise stem reserves is regarded as a beneficial trait for wheat productivity under stress (Bidinger et al., 1977; Blum, 1998; Rebetzke et al., 2008). At 12 DAA, WSC content in the stem is at its peak (Zhang et al., 2015; Shirdelmoghanloo et al., 2016). Our results showed that the stem was the main storage organ for WSC in tall genotypes in which the stem was the largest organ by weight, but not in more modern varieties where stems are much shorter as a consequence of the introduction of semi-dwarfing *Rht* genes (Borrell et al., 1993). More recent varieties partitioned more of the biomass to spikes, and the reproductive organ was also the major store of WSC in modern varieties (Fig. 4). Interestingly, in our experiment, a positive relationship between WSC concentration in the stem and total grain weight in the main spike at harvest was only observed in modern genotypes, which suggests that the important contribution of WSC content stored in the stem to grain filling may be a consequence of the introduction of semi-dwarfing genes (Richards, 1992; Miralles et al., 1998). Alternatively, it might suggest that plant breeders have selected for varieties that partition more of their WSC to spike tissues in the hot and dry conditions of South Eastern Australia, the origin of the more modern varieties. Older genotypes had large reserves of WSC in the stem for a limited sink in the spike (lower grain number), which could explain the absence of a relationship between both traits as stored WSC in the stem may not have been used (Borrell et al., 1993). In contrast with older types, more modern varieties had relatively low WSC concentration in the stem, indicating an opportunity to increase stem capacity for WSC storage in modern varieties.

### WSC availability in grains rather than grain capacity limited grain weight under stress

Starch is the main component of final grain mass. It is synthesised from stored WSC or produced from current photosynthesis. Drought and heat stress alter WSC supply to grains, either by limiting carbon assimilation through photosynthesis or by interrupting assimilate remobilisation, thus WSC availability in grains might be a limiting factor for starch synthesis and grain filling (Jurgens et al., 1978). In a field study on wheat genotypes grown in well-watered conditions, Fahy et al. (2018) quantified WSC and starch content, and key starch biosynthesis enzyme activity in grains at different grain developmental stages. They did not find any correlation between carbohydrate content in grains and final grain yield, suggesting that assimilate availability for starch synthesis is not a limiting factor for grain filling in wheat in favourable growing conditions. These findings are in accordance with our results in well-watered conditions where there was no relationship between WSC and starch content and plant yield. However, WSC concentration was reduced with increased stress intensity, and grains with relatively higher WSC concentrations at 12 DAA had higher yield (Fig. 5c). Glucose and fructose are the first substrates in the starch biosynthesis pathway (Emes et al., 2003). Genotypes with higher glucose and fructose concentrations in grains at 12 DAA had higher yield, implying that shortage in glucose and fructose might have limited starch biosynthesis later during grain filling and consequently final grain weight. Accelerated starch biosynthesis under stress depleted glucose and fructose in grains without any increase in sucrose content, indicating that insufficient sucrose supply to grains probably limited starch biosynthesis under stress.

Many studies propose sink strength (grain capacity) is the limiting factor for starch accumulation and grain filling in favourable environments (Borrás et al., 2004; Borrill et al., 2015; Fahy et al., 2018). In our study, genotypes with higher grain capacity, represented by grain dry weight at 12 DAA, had a higher yield in the well-watered treatment (Fig. 1a-b) suggesting that grain sink strength was a major determinant of grain weight at harvest when conditions were favourable. With D&H stress, however, genotypes with higher grain capacity did not have higher yield, suggesting that high grain capacity was not sufficient to determine grain weight at harvest under stress, as has also been observed in barley (Savin and Nicolas, 1996). In our experiments, drought and combined D&H did not immediately reduce grain weight at 12 DAA. Reduction in grain filling occurred after 12 DAA, which corresponds to the end of cell enlargement and beginning of carbohydrate accumulation (Emes et al., 2003). Reduced grain weight under stress was due to altered grain filling, probably as a consequence of limited WSC supply to grains.

## Conclusions

Drought and heat stress have rarely been studied together, despite their co-occurrence being a common scenario in wheat-growing regions. This work illustrated the effect of morphological changes introduced in wheat over a century on plant water use and carbohydrates partitioning. Results showed that heat stress occurring during grain filling, while plants were suffering from water stress, changed subsequent water use immediately so that some genotypes were unable to recover. Sensitivity to increased stress intensity was associated with low transpiration response to high VPD following heat stress and to genetic variation in transpiration. Reduced availability of WSC in grains following combined D&H was also identified and important for final grain weight. This suggested that measurements of transpiration and WSC content in grains following heat stress might be used to identify genetic variation for tolerance of combined drought and heat stress.

## Material and methods

### Experiment 1

#### Genetic material and growth conditions

Eight bread wheat (*T. aestivum* L.) genotypes were selected from a diverse panel of 534 wheat accessions from 44 countries described in Garcia et al. (2019). The diversity panel was previously subjected to post-anthesis drought and combined drought and heat stress in a pilot experiment and evaluated for plant total grain weight (yield) at harvest (data not shown). The selected genotypes contrasted for grain weight following drought or combined drought and heat stress, and consisted of three Australian older varieties (Currawa, Koda, Mendos), three Australian modern commercial varieties (Frame, Young, Gladius), one synthetic line from CIMMYT (Synthetic W7984) and one landrace from Ethiopia (Odessa ES19565) (Supp. Table S2). The selected genotypes were released between 1912 and 2007. In this study, Frame, Gladius and Young were considered as modern genotypes; the remaining genotypes were considered as older genotypes.

Single seeds were sown in 40 cm x 15 cm round pots containing 8.2 kg of a mixture of 1:1:1 (v/v/v) clay/loam:UC Davis mix:cocopeat mix. Seeds were sown on 11 August 2016, late winter in the southern hemisphere. From 13 days after sowing (DAS) until the end of the experiment, plants were grown in a glasshouse (34°58’17.8”S, 138°38’23.4”E) on a gravimetric platform (Droughtspotter, Phenospex, Heerlen, The Netherlands) that automatically irrigated to the pre-defined pot weight and recorded weights and water added (details in *Water use and transpiration* below). The 168 pots were randomized to 168 Droughtspotter cells using a factorial, randomized complete block design, such that each block comprised one replicate of each Genotype–Treatment combination, except in three blocks that contained one empty pot each to estimate soil evaporation. The three treatment groups comprised well-watered (WW), drought (D) and combined drought & heat stress (D&H). In particular, all plants were well-watered (soil water potential = - 0.3 MPa, gravimetric soil water content = 20 % (g/g)) and grown in temperate conditions (22 °C/15 °C day/night) until anthesis of the main spike. Anthesis date was the first day anthers were observed on the main spike. One third of the plants (WW) were maintained in well-watered, cool conditions until harvest. The remaining plants (D, D&H) were subject to a 6d drought treatment (soil water potential = - 0.6 MPa, gravimetric soil water content = 12 % (g/g)) starting 3d after anthesis on the main spike of each individual; this was followed, in half of these plants (D&H), by a 3d heat treatment at 37 °C/ 27 °C day/ night (n=7 for each accession in each treatment). Heat treatment was imposed in an adjacent glasshouse where plants were watered to weight manually. Drought was maintained until harvest in the D and D&H groups.

LED lights (400 µE/m^2^/s) were installed above plants to minimize variations due to light intensity. A graphical representation of the experimental design is shown in Supp. Fig. S5. Environmental data are shown in Supp. Fig. S6.

#### Water use and transpiration

The gravimetric platform was configured to weigh each pot at regular time intervals. All weight and water values were automatically logged and water usage estimated hourly for each pot throughout the experiment. During the heat treatment in an adjacent glasshouse, plants were watered to weight manually at similar times as the drought treatment and weights recorded. Pots were watered at least six times daily (6am, 10am, 12pm, 2pm, 4pm and 10pm). Pots containing soil only were weighed to estimate non-transpirational water loss under WW, D and D&H treatments. The water usage is a combination of plant transpiration and evaporation from the soil surface, which was negligible in all treatments as estimated from pots containing soil only.

#### Carbohydrates quantification

The main stem and spike of three plants per genotype per treatment were sampled 12d after anthesis (DAA), i.e. one day after heat treatment in drought and heat stressed plants, and stored at -80°C for further analysis. Measurements were conducted separately on the stem, flag leaf sheath, covered peduncle, exposed peduncle, rachis, grains, palea, lemma, awns and glumes. Dry weight was obtained by weighing the samples after freeze-drying. Total WSC in each tissue were determined using the anthrone method (Yemm and Willis, 1954) with some modifications: soluble sugars were extracted with 80 % ethanol at 80 °C for 1h, then extracted with distilled water at 60 °C for 1h. The extraction was repeated as many times as needed until no coloration was observed. Supernatants were combined in the same tube for colorimetric assay. Starch content in grains was measured using the Megazyme Total Starch HK (K-TSHK 08/18, Megazyme, Bray, Ireland) according to the manufacturer’s instructions. Individual WSC measurements in grains were performed in four genotypes (Frame, Odessa, Synthetic and Young). As plant morphology and grain number varied greatly between the genotypes, WSC and starch contents were expressed as g/g DW to allow for comparison between genotypes. Glucose, fructose and sucrose were analyzed in the same samples used for total WSC analysis using high performance anion exchange chromatography with pulsed amperometric detection HPAEC-PAD (Dionex ICS-5000; Thermo Fisher Scientific, Sunnyvale, USA). Separations were performed at 30 °C and the flow rate was 0.5 mL/min. A 25 μL sample was injected on a Guard CarboPac PA20 (3 × 30mm) in series with an analytical CarboPac PA20 (3 × 150mm). The elution program consisted of 0.1M NaOH from 0 to 2 min, followed by increasing 1M sodium acetate concentration up to 20 % from 2 min to 35 min, followed by increasing 1M sodium acetate concentration up to 100 % from 35 min to 36.5 min, a steady concentration from 36.5 min to 37.5 min, followed by a 0.1M NaOH wash until return to equilibrium.

Glucose, fructose and sucrose were identified based on glucose, fructose and sucrose standards. Fructans were identified by acid hydrolysis. Two WSC samples from the stem and awns were incubated with 0.2M trifluoroacetic acid (TFA) at 80 °C for 30 min together with untreated samples. Treated and untreated samples were analyzed using HPAEC-PAD as described above. Glucose, fructose and sucrose were quantified using external standards and peak areas determined using the instrument’s Chromeleon software. Fructans were quantified using peak areas.

#### Harvest data at maturity

Four plants per genotype per treatment were harvested at maturity to measure grain yield components. Total grain weight was determined for the main spike and for the whole plant. Seed number was counted using an automatic seed counter (Contador, Pfeuffer GmbH, Germany). Biomass weight included tillers, leaves and spikes but excluded grains. Plant height was measured from the base of the main stem to the top of the highest spike excluding awns. Biomass water use efficiency (bWUE) was calculated as the ratio of total aboveground biomass to total water use per plant. Grain water use efficiency (gWUE) was calculated as the ratio of total grain weight to total water use per plant.

### Experiment 2

In order to test the reproducibility of water use results, the 2016 experiment described above was replicated in an independent experiment with four genotypes (Currawa, Synthetic W7984, Mendos and Young) in 2017 with the same settings used for plant growth and treatments, except that plants were sown one month earlier. Plant water use was recorded using the gravimetric platform and three plants per genotype and per treatment were harvested at maturity and measured as before.

### Statistical analyses of yield components

The data were analysed by two-way ANOVA with genotype and treatment as fixed factors for all measured yield component and biomass traits and for the analysis of carbohydrates within each tissue. Treatment means within genotypes were compared using Tukey’s HSD (honestly significant difference) test at p = <0.1. Statistical analyses (ANOVA, Tukey’s tests, correlation analyses) and graphical representation were performed using R software (version 3.4.4, R Core Team, 2019) and ASReml-R (Butler et al., 2009).

### Statistical analyses of water use and transpiration

The recorded water use data were used to identify genotype and treatment effects on hourly transpiration rate (TR, mL/hr) and specific transpiration rate (STR, mL/hr/g biomass), with the proviso that soil evaporation and plant transpiration water losses were indistinguishable.

VPD was computed hourly from vapour capacity (VC, kPa) using the following formula:

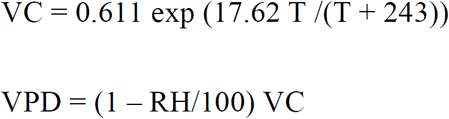

where T is temperature and RH is relative humidity. It was further decided to model TR or STR (henceforth denoted *y*) as a simple linear function of VPD with (a) genotype × treatment interaction incorporated into the VPD slope and intercept parameters, and (b) error variance modelled as a function of treatment (but not genotype). The resulting model comprises 2 x 8 x 3 = 48 fixed effects of interest (i.e. slope and intercept parameters) and 3 variance estimates, represented symbolically as

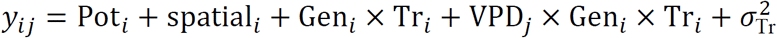

where (a) *y*_*ij*_ is the TR or STR value for Pot *i* on hour *j*, (b) Pot_*i*_ is a random-effects term for variation between pots, and (c) *spatial* comprises fixed-effects terms for spatial variation within the greenhouse. The model was fitted separately to each of TR and STR using the R package ASReml-R4 (Butler et al., 2017). Analysis of the effect of 3d heat stress on transpiration was narrowed to 30d (0 to 30 days after treatment – DAT) after heat stress, while all genotypes were still using water in the well-watered treatment, to limit the effects of intrinsic differences of grain-filling duration on transpiration and distinguish the effects of treatments.

## Supplemental data

**Supplementary figure S1.** Relationship between total water used and final grain weight per plant during Experiment 2 (2017).

**Supplementary figure S2.** Water use per plant during the three-day heat treatment, normalised to aboveground biomass.

**Supplementary figure S3.** Relationship between aboveground biomass (excluding grains) and total water used.

**Supplementary figure S4.** Genotypic differences in final grain weight (a), transpiration rate at VPD = 2 kPa (b), plant height (c), harvest index (d), grain water use efficiency (e) and WSC concentration in the stem (f).

**Supplementary table S1.** Statistical significance of differences in percentages of water-soluble carbohydrates in different parts of wheat plants: stem, grains and spike, between WW and D, D and D&H, or WW and D&H.

**Supplementary table S2.** Origins and pedigrees of the eight wheat genotypes used in the study.

**Supplementary figure S5.** (a) Schematic of the treatment design. (b) Images of plants of different genotypes following combined D&H stress at 12 DAA.

**Supplementary figure S6.** Daily maximum and minimum temperature (a), maximum daily light intensity (b), and daily maximum and minimum VPD (c) in the glasshouse experiments in 2016 and 2017.

## Acknowledgments

We thank and acknowledge the use of the facilities and scientific and technical assistance of the Australian Plant Phenomics Facility, which is supported by the Australian Government’s National Collaborative Research Infrastructure Strategy (NCRIS). In particular, we thank Chris Brien of the APPF fo his constructive comments and advise on analysis of these data. We thank Priyanka Kalambettu, Coline de l’Hommeau and Pauline Perrodin for technical assistance. We acknowledge and thank Vincent Bulone and Jelle Lahnstein for helpful comments and assistance with the carbohydrates analysis.

## Notes

Funding Information: This work was supported by the Australian Research Council Industrial Transformation Research Hub for wheat in a hot and dry climate (IH130200027).

